# APPTEST is an innovative new method for the automatic prediction of peptide tertiary structures

**DOI:** 10.1101/2021.03.09.434600

**Authors:** Patrick Brendan Timmons, Chandralal M. Hewage

## Abstract

Good knowledge of a peptide’s tertiary structure is important for understanding its function and its interactions with its biological targets. APPTEST is a novel computational method that employs a neural network architecture and simulated annealing methods for the prediction of peptide tertiary structure from the primary sequence. APPTEST works for both linear and cyclic peptides of 5-40 natural amino acids. APPTEST is computationally efficient, returning predicted structures within a number of minutes. APPTEST performance was evaluated on a set of 356 test peptides; the best structure predicted for each peptide deviated by an average of 1.9Å from its experimentally determined backbone conformation, and a native or near-native structure was predicted for 97% of the target sequences. A comparison of APPTEST performance with PEP-FOLD, PEPstrMOD and Peplook across benchmark datasets of short, long and cyclic peptides shows that on average APPTEST produces structures more-native than the existing methods in all three categories. This innovative, cutting-edge peptide structure prediction method is available as an online web server at https://research.timmons.eu/apptest, facilitating *in silico* study and design of peptides by the wider research community.

## 1 Introduction

Interest in peptide therapeutics has grown significantly in recent years, motivated in part by the many advantages peptides possess compared to traditional small molecule chemical drugs^[1,2]^. Peptide therapeutics are more selective, specific and efficacious than small molecule drugs, and are degraded to amino acids, which are less likely to exhibit undesirable drug-drug interactions. Furthermore, peptides are less likely to accumulate in tissues due to their short half-life, are less susceptible to the development of drug resistance, and are cost-effective to produce^[3–6]^.

Peptides are an important and ancient element of the immune response of all life forms, with host defence peptides (HDPs) having been identified in all species, and found to be highly conserved in vertebrate, insect and plant genomes.^[7,8]^ Peptide therapeutics are typically short, with a sequence length of 5-30 amino acids, and were initially isolated from plant or animal secretions, but are now also obtained from chemical^[9]^, genetic^[10]^ and recombinant^[11]^ libraries. Combined, these libraries present a largely unexplored chemical space. Peptides have been identified with antibacterial, antifungal, antiparasitic, antiviral and anticancer properties. A limited number of peptide drugs are currently available for treatment, including Bacitracin, Boceprevir, Enfuvirtide and Leuprolide, for treatment of pneumonia, hepatitis-C, HIV and prostate cancer, respectively. Therapeutic peptides have been found to possess *α*-helical, *β*-sheet and extended conformations in the presence of membrane or membranomimetic environments^[12–17]^.

The quantity of sequence data available from sequencing experiments has grown rapidly in recent years. The quantity of sequences with experimentally determined tertiary structures, however, is lagging behind, as determining structures experimentally is a cost and time-intensive task. Given that peptides’ bioactivities are dependent on their structure, being able to easily obtain the peptides’ tertiary structure from their primary sequence would facilitate an acceleration of the peptide drug design pipeline.

The prediction of protein structures from their primary sequences represents one of the most challenging problems in bioinformatics today. Many attempts have been made at solving the protein structure prediction problem, with many software applications having been developed for this purpose, including I-TASSER^[18]^, Rosetta^[19]^, HHpred^[20]^, NovaFold, and most notably AlphaFold 2 which recently performed excellently in the CASP 14 experiment. The prediction of peptide structures, which are distinguished from proteins by their short sequence length, present a similar challenge, which has not received the same attention as the former, with only a limited number of programs developed for the purpose of predicting peptide tertiary structure. While programs like AlphaFold 2 can utilise co-evolutionary information to predict interresidue contacts, the same is not always possible for peptides. A number of attempts have been made in the past at accurate prediction of peptide tertiary structure. Geocore, an *ab initio* filtering algorithm, was developed for finding native-like structures in small ensembles of conformations^[21]^. Later, PEPstr was developed for the prediction of peptide tertiary structures from predicted beta-turn and secondary structure information^[22]^. PEPstr has since been superseded by PEPstrMOD, which expands the scope to include cyclic peptides and peptides with non-natural residues^[23]^. Nicosia and Stracquadanio employed a generalized pattern search (Gps) algorithm, which uses search and poll to find peptide conformation global energy minima^[24]^. PepLook explores the peptide conformational space using a Boltzman-stochastic algorithm^[25–27]^. At a similar time, Maupetit et al. developed PEP-FOLD, which has been updated multiple times^[28–32]^. The most recent version combines a structural alphabet with a Hidden Markov model. Finally, Narzisi et al. employed a multi-objective evolutionary algorithm for the exploration of the peptide conformational space^[33]^.

Machine learning techniques, including deep learning, have previously been applied to other bioinformatic problems: DeepPPISP for the prediction of protein-protein interaction sites^[34]^, SCLpred and SCLpred-EMS for protein subcellular localization prediction^[35,36]^, CPPpred for the prediction of cell-penetrating peptides^[37]^, HAPPENN for the prediction of peptide hemolytic activity^[38]^, ENNAACT for the prediction of peptide anticancer activity,^[39]^ and ENNAVIA for the prediction of peptide antiviral activity^[**?**]^. Indeed, deep learning has been applied to the prediction of protein secondary structures^[40–43]^. Herein we describe APPTEST, a novel method for the automatic prediction of peptide tertiary structures. APPTEST utilises one-dimensional gated residual convolutional neural networks for the prediction of distance and dihedral angle restraints, which are then input to traditional NMR structure determination methods to obtain a final ensemble of model structures.

## 2 Methods

### 2.1 Dataset

The proper construction of a reliable dataset is an important step in a machine learning endeavour. Neural networks’ performance scales with the size of the dataset available for training; it is important, therefore, to construct a dataset encompassing as many peptide structures as possible from multiple sources. PDB structure codes were sourced from multiple peptide databases: DBAASP^[44,45]^, APD3 ^[46]^, ADAM^[47]^ and DRAMP^[48]^. PDB structure codes were also extracted from the datasets of PEPstrMOD^[23]^ and PEP-2D^[49]^. Finally, the RCSB PDB^[50]^ was also searched for structures with a chain length between 5-40 amino acids.

Only structures with a sequence length between 5-40 were considered. To prevent the classifier from overfitting to the training data, the dataset’s sequences were internally redundancy reduced using CD-HIT^[51–53]^, with a sequence identity cut-off of 0.9. NMR structures with an internal backbone RMSD greater than 2.50 Å were excluded.

### 2.2 Model validation

It is critically important to thoroughly validate machine learning models. Tenfold cross-validation and validation by an external test set were employed to evaluate the performance of APPTEST. 2265 experimentally obtained peptide structures were used for model training and internal tenfold cross-validation. The models trained under cross-validation were ensembled and evaluated with the external test set, which consists of 356 previously unseen, redundancy reduced peptide sequences and their corresponding experimentally obtained peptide structures.

### 2.3 Structure analysis

The Biopython^[54]^ module Bio.PDB^[55]^ was used to retrieve peptide structures from the PDB, calculate inter-residue distances, dihedral angles and root mean square deviation (RMSD) values.

### 2.4 Peptide representation

186 amino acid scales were extracted from the AAindex^[56]^, and used to construct the matrix *A*, of shape (21, 186), where the first twenty rows correspond to the twenty proteinogenic amino acids, the last row corresponds to the non-natural amino acids without known AAindex-scaled properties, and the 186 columns correspond to the 186 amino acid indices. *A*^*T*^ is scaled using the standard scaler, and dimensionality reduced with principal component analysis^[57]^ and scaled so its values have minimum and maximum values of zero and one, respectively. The final matrix *A* has shape (21, 15).

Each peptide can be represented by one-hot encoding its primary sequence to give a vector *h*, where 0 represents an empty position, 1-20 encode the natural, proteinogenic amino acids, and 22 encodes a non-natural amino acid residue. Similarly, the one-hot encoding can be represented by a sparse matrix *P* of shape (50,21) exists, where *P*_*ij*_ = 1 if the amino acid at sequence position *i* is the *j*^*th*^ of the 21 types of amino acid. Information about cyclic restraints can be encoded in a sparse matrix *C* of shape (50,50), where *C*_*i,j*_ = 1 if a cyclization exists between the *i*^*th*^ and *j*^*th*^ residues. Finally, a matrix *S*, of shape (50,15) which describes the peptide’s amino acids’ physicochemical properties can be defined as:

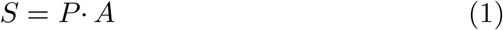

Additionally, two masks can be defined. A vector of length 50, *m*, and a matrix of shape (50,50) *M*. *m*_*i*_ = 1 if the *i*^*th*^ position of the peptide sequence space is occupied. Similarly *M*_*i,j*_ = 1 if the *i*^*th*^ and *j*^*th*^ positions of the peptide sequence space are occupied.

### 2.5 Data augmentation

In order to artificially increase the quantity of data available, data augmentation is employed. Specifically, the peptides’ inputs to the neural networks are shifted along the input frame of length 50. During training, each peptide is represented as 20 samples, each randomly shifted along the input frame. During blind prediction, each peptide is represented as *n* samples, where *n* is 50-(peptide length)+1. The neural networks’ outputs are shifted back to the original frame, and averaged.

### 2.6 Neural network architecture and implementation

Keras with a Tensorflow^[58]^ backend was used for the construction and training of the neural network. A randomized grid search strategy was employed for the identification of the optimal neural network architecture and hyperparameters.

First, *h* is input to an embedding layer with a dense embedding dimension of 12. Each row of the dense embedding is then multiplied by the mask vector *m*, resulting in a final tensor of shape (50,15). This tensor is concatenated with *A* and *C* to yield a final tensor of shape (50, 77), which is input to a one-dimensional convolutional layer, with 128 filters and a window width of 7. This is followed by batch normalization layer, the rectified linear unit activation function, and a one-dimensional spatial dropout layer, and two residual gated convolutional blocks.

Each residual gated convolutional block consists of three one-dimensional gated convolutional layers^[59]^, which also have 128 filters and a window width of 7. The first two are followed by a batch normalization layer and a rectified linear unit activation function, and the final has a spatial dropout applied to it. The output of the spatial dropout layer is added to the block’s original input, batch normalized, activated with the rectified linear unit and has another spatial dropout layer applied. The output of the second residual gated convolutional block is connected to a fully connected layer with 1024 nodes, which is followed by a batch normalization, a rectified linear unit activation and a dropout layer. This layer is connected to three output layers, which have 2500, 2500 and 200 nodes, respectively. The first two are activated with the rectified linear unit, and the third is activated with the hyperbolic tangent function. When reshaped, and multiplied with their respective masks, these output layers correspond to the C_*α*_-C_*α*_ and C_*β*_-C_*β*_ distances, and the cos and sin of the peptide’s *φ* and *ψ* dihedral angles.

The mean squared error function (MSE), commonly used for regression tasks, is employed as the loss function in training the neural network. It is defined as:

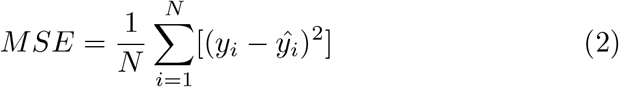

where *y*_*i*_ is the true value of the *i*^*th*^ sample, and 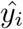 is the predicted value of the *i*^*th*^ sample.

Adaptive Momentum with Nesterov momentum (Nadam) was identified as the optimal optimizer^[60]^.

The neural network was trained for 400 epochs, without stopping criteria. The model with the lowest validation MSE encountered during training was retained for each of the cross-validation splits. The optimal learning rate parameter was found to be 0.001.

### 2.7 Simulated annealing protocols

#### 2.7.1 Distance restraints

*C*_*α*_ − *C*_*α*_ and *C*_*β*_ − *C*_*β*_ distance restraints are derived from the neural networks’ predictions, with the lower distance restraints being calculated as *mean* − *sd*, and the upper distance restraints being calculated as *mean* + *sd*.

#### 2.7.2 Dihedral restraints

The predicted values for each dihedral angle’s cos and sin values are averaged, and those average values are used to recover a predicted dihedral angle value. The upper and lower dihedral angle restraints are given as *mean* + 15° and *mean* − 15°, respectively. Only predictions where both the cos and sin standard deviations are below 0.10 are included as distance restraints.

#### 2.7.3. XPLOR-NIH protocol

XPLOR-NIH 3.1 is used for simulated annealing and energy minimization, for a default of 100 structures^[61,62]^. Structure coordinates are initiated from a preliminary structure constructed using Peptide-Builder^[63]^, based on neural networks’ predicted dihedral angles. Upper and lower distance and torsion angle restraints are loaded, and torsion angle dynamics are performed, with an initial temperature of 2025 K and a final temperature of 25 K. A temperature step of 25 K is employed, with a tolerance factor of 100 for the annealing stage. Finally, cartesian angle minimization is performed.

#### 2.7.4 CYANA protocol

Alternatively, CYANA 3.0 is also used for simulated annealing^[64]^. The upper and lower distance and torsion angle restraints are loaded, and by default, 100 random structures are created, are annealed for 20,000 steps and energy minimized for 20,000 steps.

### 2.8 Performance evaluation

The robustness of the predictions is evaluated by measurement of the backbone root mean square deviation (B-RMSD) between the predicted and experimental structures. Results are reported for both the best-predicted model, and the ensemble’s prime model. The choice of the ensemble’s prime model is dependent on the modelling software employed: CYANA models are ordered by target function value, while XPLOR-NIH models are by default ordered by total energy. B-RMSD values are calculated for both the full structure, and the rigid core. A structure is considered near-native, if the rigid-core B-RMSD to the experimental conformation is *<*4Å, as previously defined by Thévenet et al. ^[30]^.

### 2.9 Rigid core

As NMR model structures can exhibit significant structural diversity, ensemble-level comparisons between the predicted and experimental structures are performed for the peptides’ rigid cores as well as the full structures. Rigid core (RC) regions were calculated using the method described by Maupetit et al. ^[29]^ A peptide’s rigid core is defined as the set of residues that exhibit a C*α*-RMSD of *<*1.5 Å.

### 2.10 Comparison with existing methods

The predictive performance of APPTEST is compared with the existing peptide tertiary structure prediction methods. The structures selected for the comparison are those used for similar comparison tables in the articles describing PEPstr^[22]^, PEPstrMOD^[23]^, PEP-FOLD^[28–31]^ and Peplook^[27]^.

APPTEST is used with XPLOR-NIH to predict 100 structures. The structures are sorted by their total energies. PEPFOLD 3.5 is used to predict 100 structures. The PEP-FOLD structures are sorted by the sOPEP energy, and all 100 structures are retained. For cyclic peptides, PEPFOLD 2.0 is used, as recommended by the authors, with the recommended short simulation time employed. PEPstrMOD only predicts a single structure for each query. The simulation time used is 100 ps in a vacuum environment. Peplook does not have an online interface that can be used for structure prediction; consequently, it was not possible to independently evaluate the Peplook method. The results presented are reproduced directly from Beaufays et al.^[27]^.

## 3 Results and Discussion

### 3.1 Model validation

In order to comprehensively evaluate the predictive ability of APPTEST, predicted structures were calculated for the 356 peptide sequences of the independent test set. The full set of results is detailed in Supplementary Table 1. A summary of the structural statistics is given in Table 1.

**Table 1:** APPTEST performances on the APPTEST independent test set, using CYANA and XPLOR-NIH for torsion angle dynamics and simulated annealing. B-RMSD values are given for the best-predicted model, and the model with the least distance restraint violations, and the model with the lowest energy (XPLOR-NIH only). Numbers in brackets are B-RMSD values for the peptides’ rigid cores only.

| Program |  | CYANA |  | XPLOR-NIH |  |  |
| --- | --- | --- | --- | --- | --- | --- |
| LENGTH | N | BEST | RESTR. | BEST | ENERGY | RESTR. |
| 6-12 | 123 | 1.85 (1.83) | 2.43 (2.41) | 1.57 (1.56) | 2.30 (2.28) | 2.45 (2.44) |
| 13-19 | 83 | 2.07 (1.95) | 2.63 (2.47) | 1.93 (1.80) | 2.70 (2.52) | 2.71 (2.56) |
| 20-26 | 44 | 2.57 (2.33) | 3.25 (2.96) | 2.39 (2.18) | 3.05 (2.74) | 3.12 (2.83) |
| 27-33 | 52 | 2.21 (2.01) | 2.71 (2.48) | 2.07 (1.88) | 2.69 (2.42) | 2.65 (2.41) |
| 34-40 | 41 | 2.59 (2.30) | 3.00 (2.69) | 2.50 (2.20) | 3.07 (2.69) | 3.36 (3.00) |
| All | 356 | 2.10 (1.98) | 2.65 (2.51) | 1.91 (1.79) | 2.61 (2.44) | 2.70 (2.54) |

While APPTEST itself generates distance and dihedral restraints, a molecular modelling program is required for the creation of model structures based on these restraints. Two molecular modelling packages are employed and compared in this work: CYANA and XPLOR-NIH. The former benefits from faster computation times, but requires a license for use, which precludes us from integrating it with our web server. The web server can create the required CYANA-format restraints which the user can use for the creation of structures with their local copy of CYANA or any server that supports CYANA. The latter, while computationally slower, can be downloaded with a license for free for academic use from the authors’ website.

The structure prediction results tabulated for the 356 peptides in Table 1 show that APPTEST is a reliable method for the prediction of structures of peptides of 5-40 amino acids. The performance with both packages is comparable, although the better performance is achieved with the XPLOR-NIH package, with a mean best B-RMSD of 1.91 Å, compared to a B-RMSD of 2.10 Å when using CYANA for the 356 peptides. A mean B-RMSD of just 1.57 Å is achieved for short peptides between 6-12 amino acids in length, rising to just 2.50 Å for longer peptides with between 34-40 amino acids. The best RMSD achieved was 0.23 Å for the structure 2mjr, which is 10 amino acids long, while the worst RMSD was 8.38 Å for the structure 2ki0. An RMSD below 3.00 Å is achieved for 84% of the 356 structures tested, and an RMSD below 2.00 Å is achieved for 60% of the structures tested. Furthermore, only 3% of peptides have a best B-RMSD greater than 4.00 Å.

**Figure 1.**
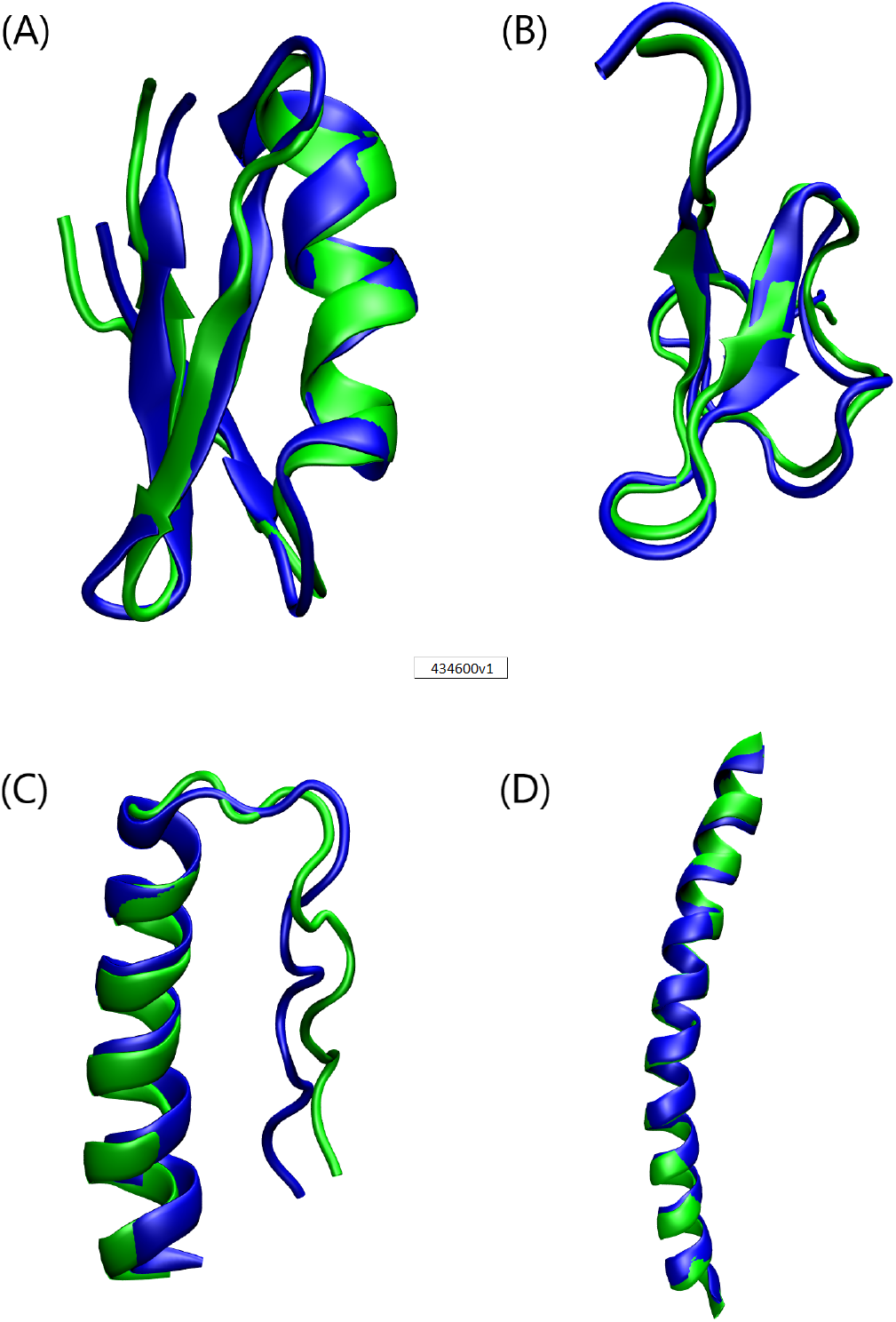
Selection of some of the best APPTEST predicted structures (green) aligned with the peptides’ corresponding experimental conformations (green). (A) Hongotoxin 1 (1hly) is a 39 amino acid mixed-structure peptide, which has both *α*-helical and *β*-sheet structures^[65]^. The best APPTEST structure has a B-RMSD of 1.44 (1.38) Å to the experimental conformation. (B) Jingzhaotoxin-XI (2a2v) is a 34 residue peptide with two *β*-sheets^[66]^. The best APPTEST structure has a B-RMSD of 1.46 (1.31) Å to the solution NMR structure. (C) Optimised PPa-TYR (6gwx) is a 36 residue rationally designed miniprotein with a polyproline-II helix and an *α*-helix motif^[65]^. The best APPTEST structure has a B-RMSD of 1.75 (1.74) Å to the experimentally determined structure1.5(D) The hydrophobic analogue of winter flounder antifreeze protein (1j5b) is a 37 amino acid *α*-helical structure. The APPTEST predicted structure with the lowest energy is the best-predicted structure, with a B-RMSD of just 0.86 Å to the experimental conformation^[67]^.

### 3.2 Comparison with existing methods

To effectively benchmark the predictive performance of APPTEST against PEP-FOLD, PEPstrMOD and Peplook, the structures predicted by each method were compared to the experimental structures and the B-RMSD values were measured. Three benchmark sets of structures are used: short peptides (9-25 aa), long peptides (26-40 aa) and cyclic peptides (10-30 aa). The results are detailed in Tables 2, 3 and 4.

**Table 2:** Performance comparison of APPTEST, PEP-FOLD 3.5 and PEPstrMOD B-RMSD values on peptides with 9-25 amino acids. Numbers in brackets are B-RMSD values for the peptides’ rigid cores only.

| PDB | L | RC | APPTTEST | PEPFOLD | PEPstrMOD | APPTTEST | PEPFOLD |
| --- | --- | --- | --- | --- | --- | --- | --- |
|  |  |  | PRIME | PRIME | ONLY | BEST | BEST |
| 1a13 | 14 | 2–14 | 2.26 (2.05) | 2.05 (1.85) | 3.01 (2.98) | 2.02 (1.89) | 1.82 (1.73) |
| 1b03A | 18 | – | 5.05 | 2.69 | 4.26 | 2.85 | 2.33 |
| 1du1 | 20 | – | 4.94 | 5.12 | 5.63 | 4.16 | 4.88 |
| 1e0q | 17 | – | 3.77 | 2.31 | 4.54 | 2.79 | 0.98 |
| 1egs | 9 | – | 6.52 | 2.04 | 2.08 | 3.65 | 1.05 |
| 1gjf | 14 | 4–14 | 1.86 (0.78) | 2.38 (0.73) | 6.53 (5.28) | 0.96 (0.55) | 1.39 (0.66) |
| 1in3 | 12 | – | 1.24 | 1.67 | 2.50 | 1.20 | 1.55 |
| 1l2y | 20 | – | 1.59 | 2.90 | 6.24 | 1.26 (1.26) | 1.18 |
| 1l3q | 12 | – | 5.98 | 3.92 | 3.14 | 3.80 (3.80) | 2.57 |
| 1lcx | 13 | 3–12 | 2.25 (1.20) | 2.51 (1.46) | 4.30 (3.15) | 1.80 (1.12) | 2.42 (1.38) |
| 1niz | 14 | – | 2.32 | 1.68 | 5.55 | 1.74 | 1.22 |
| 1nkf | 16 | – | 5.41 | 4.71 | 4.26 | 4.13 | 1.56 |
| 1pef | 18 | – | 0.60 | 0.90 | 0.99 | 0.60 | 0.69 |
| 1rpv | 17 | 4–16 | 0.86 (0.35) | 0.74 (0.54) | 5.57 (5.46) | 0.63 (0.28) | 0.73 (0.38) |
| 2bta | 15 | 4–9 | 4.53 (2.61) | 4.50 (2.60) | 4.17 (3.26) | 3.08 (1.70) | 4.36 (2.51) |
| 1c98 | 10 | – | 2.84 | 5.39 | 4.63 | 2.30 | 2.60 |
| 1d6x | 13 | 1–12 | 2.53 (2.50) | 2.94 (2.94) | 4.59 (4.51) | 2.34 (2.16) | 2.29 (2.20) |
| 1d7n | 14 | 2–13 | 1.08 (0.69) | 1.04 (0.64) | 2.88 (2.23) | 0.89 (0.61) | 0.97 (0.62) |
| 1d9j | 20 | 10–20 | 2.06 (0.89) | 4.38 (1.18) | 6.12 (4.03) | 1.50 (0.61) | 2.58 (0.98) |
| 1d9l | 17 | 4–15 | 1.47 (0.47) | 1.71 (0.67) | 5.83 (4.93) | 1.25 (0.39) | 1.67 (0.62) |
| 1d9m | 18 | 10–18 | 1.95 (1.31) | 2.62 (1.16) | 4.38 (3.46) | 1.66 (0.58) | 2.27 (0.97) |
| 1d9o | 20 | 11–20 | 3.17 (0.44) | 3.10 (0.51) | 5.23 (3.17) | 2.47 (0.34) | 2.78 (0.43) |
| 1d9p | 20 | 10–20 | 1.88 (0.54) | 2.18 (0.63) | 3.52 (3.33) | 1.62 (0.46) | 1.92 (0.57) |
| 1dn3 | 15 | – | 1.10 | 1.14 | 7.20 | 1.02 ( | 0.91 |
| 1g89 | 13 | 1–12 | 2.55 (2.34) | 4.51 (4.36) | 3.15 (3.13) | 1.82 (1.75) | 3.28 (3.15) |
| 1hu5 | 18 | 2–18 | 1.96 (1.79) | 1.60 (1.54) | 2.49 (2.45) | 1.61 (1.48) | 1.39 (1.33) |
| 1hu6 | 18 | 2–18 | 3.26 (3.25) | 3.43 (3.43) | 4.26 (4.09) | 2.51 (2.50) | 2.71 (2.43) |
| 1hu7 | 18 | 2–18 | 2.05 (1.94) | 1.79 (1.63) | 3.33 (3.31) | 1.77 (1.69) | 1.79 (1.58) |
| 1id6 | 15 | 1–14 | 5.54 (5.59) | 5.50 (5.55) | 7.42 (7.4) | 4.96 (5.02) | 4.50 (4.48) |
| 1jav | 19 | – | 1.45 | 6.13 | 7.44 | 1.43 | 3.28 |
| 1kzv | 18 | 2–18 | 1.20 (1.02) | 1.78 (1.65) | 4.73 (4.34) | 0.79 (0.72) | 1.57 (1.44) |
| 1m02 | 12 | 2–11 | 4.23 (3.13) | 4.30 (3.33) | 5.89 (4.30) | 2.39 (1.86) | 3.01 (2.75) |
| 1myu | 12 | – | 1.73 | 3.00 | 4.38 | 1.22 | 0.96 |
| 1odp | 20 | – | 1.83 | 1.89 | 4.75 | 1.34 | 1.75 |
| 1p0j | 19 | 2–19 | 1.58 (1.30) | 1.72 (1.57) | 4.04 (3.92) | 1.27 (1.11) | 1.57 (1.39) |
| 1p0l | 19 | 2–19 | 1.66 (1.28) | 1.88 (1.59) | 5.12 (5.22) | 1.27 (1.15) | 1.69 (1.49) |
| 1p0o | 19 | 2–19 | 1.54 (1.34) | 1.98 (1.87) | 4.77 (4.61) | 1.29 (1.06) | 1.65 (1.52) |
| 1p5k | 19 | 2–18 | 1.29 (1.03) | 1.77 (1.54) | 4.74 (4.16) | 1.21 (0.72) | 1.62 (1.35) |
| 1qcm | 11 | 2–11 | 2.95 (2.99) | 2.49 (1.90) | 3.92 (3.71) | 2.72 (2.69) | 1.10 (1.02) |
| 1qfa | 13 | – | 1.01 | 0.72 | 5.34 | 0.90 | 0.66 |
| 1sol | 20 | – | 4.14 | 3.27 | 6.38 | 2.56 | 2.59 |
| 2bp4 | 16 | 1–15 | 2.17 (1.64) | 5.45 (5.21) | 6.31 (6.15) | 1.64 (1.60) | 4.12 (4.06) |
| MEAN |  |  | 2.60 (2.24) | 2.81 (2.38) | 4.66 (4.33) | 1.96 (1.69) | 2.05 (1.71) |

PDB: Protein Data Bank identifier. L: peptide length. RC: rigid core. B-RMSD:
Backbone root mean square deviation. Results are reported for the best model and the first model, for both the full structure and the rigid core. APPTEST first model is that with the lowest XPLOR-NIH energy. PEP-FOLD first model is the one with the lowest sOPEP energy value. PEPstrMOD only returns a single structure. Numbers within brackets are rigid core B-RMSDs, where applicable.

**Table 3:** Performance comparison of APPTEST, PEP-FOLD 3.5 and PEPstrMOD B-RMSD values on peptides with 26-40 amino acids. Numbers in brackets are B-RMSD values for the peptides’ rigid cores only.

| PDB | L | RC | APPTTEST | PEPFOLD | APPTTEST | PEPFOLD |
| --- | --- | --- | --- | --- | --- | --- |
|  |  |  | PRIME |  | BEST |  |
| 1by0 | 27 | 1–23 | 1.68 (0.98) | 1.65 (1.38) | 1.42 (0.96) | 1.37 (1.22) |
| 1yyb | 26 | 1–20 | 2.65 (1.57) | 3.26 (1.29) | 1.89 (1.03) | 1.85 (1.00) |
| 2kbl | 27 | 6–27 | 2.35 (2.43) | 7.50 (5.79) | 1.92 (1.87) | 3.35 (2.14) |
| 2k76 | 30 | 4–29 | 9.91 (6.75) | 4.40 (3.77) | 5.95 (4.46) | 1.29 (1.03) |
| 2gdl | 31 | 8,10–11,14–15,21–29 | 6.32 (4.90) | 7.36 (4.69) | 4.94 (4.07) | 5.01 (4.14) |
| 2l0g | 32 | 5–32 | 3.73 (1.82) | 3.57 (1.48) | 2.27 (1.36) | 1.70 (1.04) |
| 2bn6 | 33 | 4–29 | 2.17 (1.01) | 12.44 (9.69) | 1.62 (0.95) | 8.32 (6.42) |
| 2ovc | 30 | – | 2.89 | 10.39 | 2.4 | 7.16 |
| 1bwx | 39 | 15–28 | 7.06 (0.90) | 4.31 (1.45) | 5.24 (0.74) | 4.06 (1.42) |
| 2kya | 34 | 11–30 | 3.37 (1.25) | 8.52 (6.61) | 3.09 (0.97) | 6.74 (4.35) |
| 1wy3 | 35 | 1–26 | 1.39 (1.44) | 4.26 (3.58) | 1.35 (1.32) | 3.18 (2.60) |
| 1wr3 | 36 | 5–15,17–34 | 1.47 (1.26) | 5.01 (3.39) | 1.47 (1.26) | 2.61 (1.80) |
| 1wr4 | 36 | 5–34 | 7.36 (6.67) | 5.36 (3.67) | 2.22 (1.64) | 2.08 (1.67) |
| 2ki0 | 36 | 5–11,13–36 | 11.92 (10.82) | 2.51 (1.89) | 8.38 (8.48) | 2.18 (1.80) |
| 1e0m | 37 | 3–35 | 2.75 (2.32) | 4.16 (4.20) | 2.07 (1.79) | 2.51 (2.33) |
| 1e0n | 27 | 1–25 | 6.85 (6.42) | 6.79 (6.29) | 3.73 (3.75) | 5.10 (5.19) |
| 1yiu | 37 | 5–36 | 2.11 (1.86) | 5.67 (4.91) | 2.00 (1.61) | 2.90 (2.40) |
| 1bhi | 38 | 7–11,14–33 | 4.02 (1.74) | 7.26 (4.78) | 3.19 (1.40) | 4.44 (2.83) |
| 1i6c | 39 | 2–11,16–33 | 4.52 (2.03) | 6.98 (4.32) | 3.68 (1.92) | 3.57 (1.75) |
| 1jrj | 39 | 11–38 | 2.22 (1.31) | 3.72 (2.71) | 2.06 (1.23) | 3.06 (2.57) |
| 2ysc | 39 | 8–39 | 5.03 (4.81) | 5.96 (4.98) | 4.07 (3.89) | 3.76 (2.57) |
| 1e0l | 37 | 6–37 | 2.87 (1.83) | 5.13 (3.78) | 2.71 (1.83) | 3.61 (2.41) |
| 2ysf | 40 | 7–39 | 4.67 (3.59) | 6.98 (5.08) | 3.86 (3.56) | 3.77 (2.29) |
| 2ysg | 40 | 7–38 | 8.99 (8.32) | 6.32 (4.68) | 3.89 (3.15) | 3.42 (2.24) |
| 2ysh | 40 | 7–37 | 4.09 (2.74) | 6.18 (3.74) | 3.44 (1.99) | 3.51 (2.23) |
| 2ysi | 40 | 9–40 | 4.42 (3.52) | 8.64 (7.92) | 4.19 (3.35) | 3.77 (2.86) |
| 1ywj | 28 | – | 2.03 | 3.81 | 1.92 | 2.09 |
| 1ymz | 37 | 3–13,5–32 | 2.37 (1.48) | 5.07 (3.50) | 2.10 (1.30) | 2.57 (1.68) |
| 3e21 | 40 | – | 9.14 | 2.98 | 6.95 | 2.98 |
| 1use | 40 | – | 3.05 | 2.15 | 2.1 | 1.16 |
| MEAN |  |  | 4.45 (3.36) | 5.61 (4.30) | 3.20 (2.44) | 3.43 (2.58) |

**Table 4:**
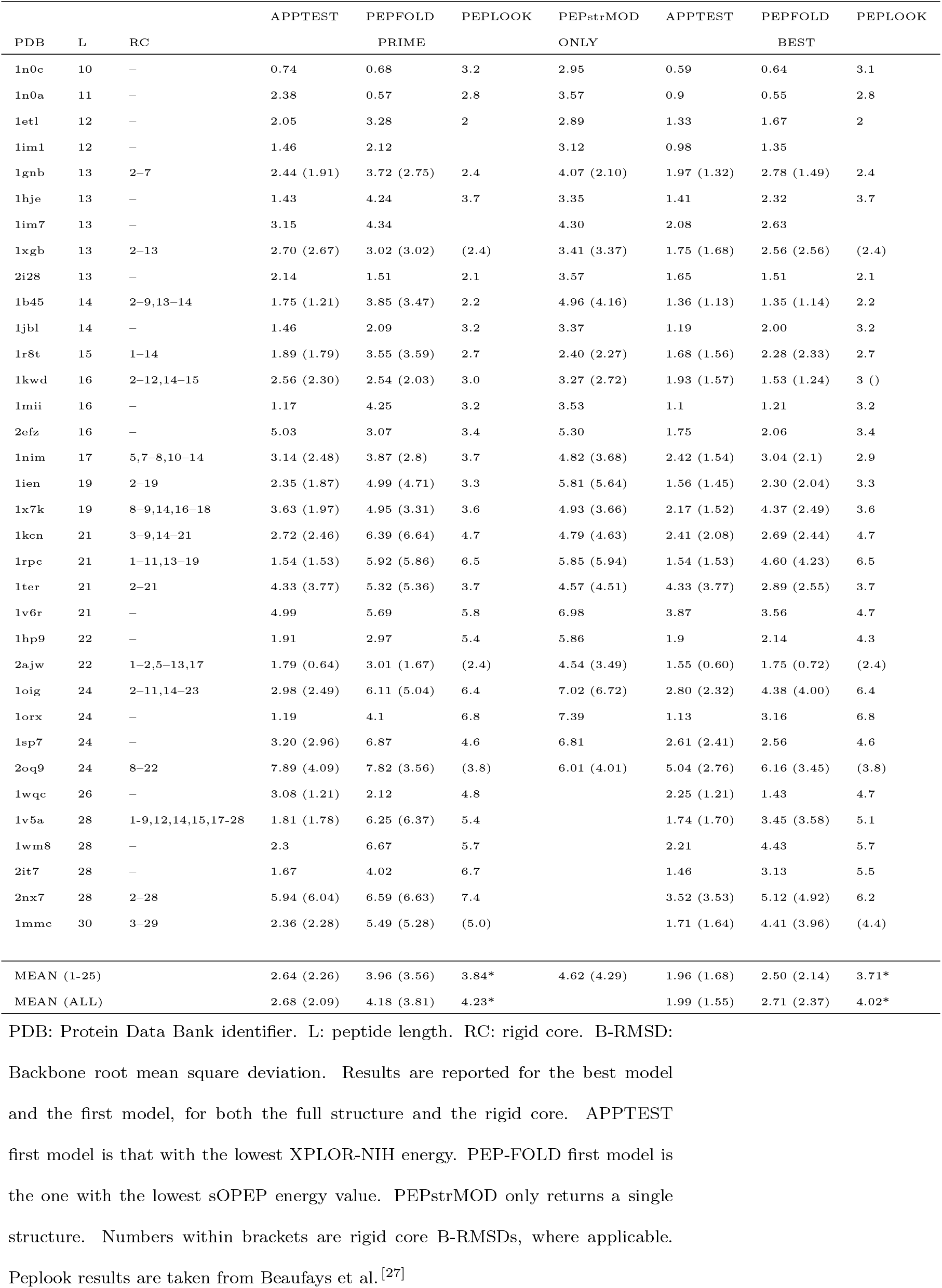
Performance comparison of APPTEST, PEP-FOLD 2.0, Peplook and PEPstrMOD B-RMSD values on cyclic peptides. Numbers in brackets are B-RMSD values for the peptides’ rigid cores only.

#### 3.2.1 Short peptides

A total of 42 short (9-25 aa) peptides are predicted. Overall, APPTEST performs best, with its mean B-RMSD values being lower than the corresponding PEP-FOLD and PEPstrMOD values. Closer inspection reveals that of the best full structure predictions, APPTEST scores best on 27 of the 42 structures, and PEP-FOLD 3.5 scores best on the remaining 15 structures. The results for just the rigid cores are similar, with APPTEST scoring best on 26 structures, and PEP-FOLD scoring best on 16 structures.

As the best structure is one of a hundred structures, the first structure, ie. the one with the lowest energy, should also be as close as possible to the experimental structure. Of the 42 structures investigated, APPTEST predicts the most-native structure for 25 of the peptides, PEP-FOLD 3.5 for 14 peptides, and PEPstrMOD for the remaining 3. The rigid core results are similar, with APPTEST, PEP-FOLD and PEPstrMOD scoring 25, 15 and 2, respectively.

#### 3.2.2 Long peptides

Prediction of peptide structures is increasingly challenging as the sequence length increases, as evidenced by APPTEST’s mean best full structure B-RMSD being 3.20 Å for long peptides (26-40 aa), compared to 1.96 Å for short peptides. Nonetheless, APPTEST still outperforms

PEP-FOLD 3.5 in this task, which has a mean B-RMSD of 3.43 Å. PEP-strMOD, meanwhile, does not facilitate predictions of peptides with a sequence length greater than 25 amino acids.

APPTEST and PEP-FOLD 3.5 return the most-native predictions for 16 and 14 of the 30 peptide structures included in this benchmark when the full structure B-RMSD are considered; this rises to 19 and 11 when only the structures’ rigid cores are considered. When the comparison is restricted to consider only the lowest energy structure returned by each method, APPTEST and PEP-FOLD 3.5 return the most-native structure for 20 and 10 of the peptides, respectively, for both the full structures and rigid core regions.

#### 3.2.3 Cyclic peptides

The structures of cyclic peptides, such as those possessing disulfide bonds, have traditionally been challenging to predict. Neither PEPstr^[22]^, the Gps algorithm^[24]^ nor PEP-FOLD 1.0 ^[28]^ facilitated the prediction of cyclic peptide structures.

The 34 cyclic peptides in this benchmark set are between 10-30 amino acids in length. As PEPstrMOD only facilitates predictions of peptides up to 25 amino acids in length, mean RMSD values are reported for the subset of 28 peptides with a sequence length of 10-25 amino acids, as well as for the entire set of 34 peptides. As the Peplook web server is no longer available, the B-RMSD values that follow are taken from the Peplook article^[27]^. Unfortunately, the values presented are for only the full structure or only the rigid core, and so the comparison with Peplook is consequently not fully comprehensive. As per the instructions on the PEP-FOLD web server, PEP-FOLD 2.0 is used instead of the newer PEP-FOLD 3.5 for the prediction of cyclic peptide structures.

Comparing the performance of the four methods on the cyclic dataset, it is clear that APPTEST outperforms the existing methods, with a mean best full structure B-RMSD value of 1.96 Å, compared to the 2.50 Å, 3.71Å and 4.62 Å of PEP-FOLD, Peplook and PEPstrMOD, respectively. This is further exemplified by inspecting predictions of individual structures: APPTEST returns the most native structure for 26 of the 34 peptides in the benchmark, PEP-FOLD 2.0 returns the most native structure for the remaining 8 peptides. Considering only the rigid core regions, APPTEST returns the most accurate structure for 30 of the 34 peptides, with PEP-FOLD 2.0 scoring better on the remaining 4 peptides. Restricting the comparison to only consider the prime structure returned by each method, APPTEST achieves the best performance on 23 of the structures, with the remaining 6, 4 and 1 structures being best predicted by PEP-FOLD 2.0, Peplook and PEPstrMOD, respectively. The results when considering only the rigid core region are similar, with APPTEST, PEP-FOLD 2.0 and Peplook achieving the best performance for 25, 7 and 2 of the 34 structures, respectively.

To conclude, an accurate method for the prediction of peptide and miniprotein tertiary structure is essential for the *in silico* design of new peptide-based therapeutics. Access to a reliable peptide structure model would allow for a reduction in the time and cost demands of the design phase. This study describes APPTEST, a novel neural method for the automated prediction of peptide and miniprotein tertiary structures. APPTEST distinguishes itself from existing peptide structure prediction methods by being based on a neural network architecture, supported by software traditionally used for the calculation of NMR solution-state structures. Our method achieves excellent performance, returning the most-native model structures for peptides, short and long, linear and cyclic. The B-RMSD values between APPTEST’s predicted structure and the experimentally determined conformations are lower than those achieved by existing methods in all categories, both for the prime and best models, whether full structure or only the rigid core. This novel method, therefore, represents a new state-of-the-art in peptide structure prediction. We hope that this work will aid future studies focused on the design of novel peptide-based therapeutics.

## 4 Availability

### 4.1 Web server

APPTEST is available as an easy to use web server online at https://research.timmons.eu/apptest, for the benefit of the wider scientific community. The web server is capable of predicting peptides’ tertiary structure based on the primary sequence and cyclic restraints. Input peptide sequences are restricted to the 20 natural amino acids; support for the prediction of peptides containing non-natural amino acids is not currently available. Users may choose to only predict the distance and angle restraints, or to also conduct simulated annealing to produce an ensemble of model structures. Simulated annealing can be carried out only using XPLOR-NIH on the web server, as the authors’ CYANA license does not extend to usage in a web server context.

### 4.2 Standalone

APPTEST is also available as a standalone executable program for Linux. This program has been tested to work with Ubuntu 20.04 LTS and Debian 10. The program can be downloaded from https://research.timmons.eu/apptest, or alternatively can be requested from the authors. The standalone program is recommended for users who intend to carry out a large number of structure predictions.

## Data Availability

Datasets employed during this study are available for download at https://research.timmons.eu/apptest

## Acknowledgments

The authors would also like to thank University College Dublin for the Research Scholarship granted to P.B.T.

